# Dual-view oblique plane microscopy (dOPM)

**DOI:** 10.1101/2020.09.24.311613

**Authors:** Hugh Sparks, Lucas Dent, Chris Bakal, Axel Behrens, Guillaume Salbreux, Chris Dunsby

## Abstract

We present a new folded dual-view oblique plane microscopy (OPM) technique termed dOPM that enables two orthogonal views of the sample to be obtained by translating a pair of tilted mirrors in refocussing space. Using a water immersion 40× 1.15 NA primary objective, deconvolved image volumes of 200 nm beads were measured to have full width at half maxima (FWHM) of 0.35±0.04 μm and 0.39±0.02 μm laterally and 0.81±0.07 μm axially. The laterally integrated z-sectioning value was 1.33±0.45 μm using light-sheet FWHM in the frames of the two views of 4.99±0.58 μm and 4.89±0.63 μm. To qualitatively demonstrate that the system can reduce shadow artefacts while providing a more isotropic resolution, a multi-cellular spheroid approximately 100 μm in diameter was imaged.

## 1. Introduction

Light-sheet fluorescence microscopy (LSFM) [1,2] provides optically sectioned fluorescence imaging with low photobleaching and photoxicity to the sample [3]. LSFM traditionally uses separate objective lenses to provide sheet-like laser illumination for fluorescence excitation and to provide orthogonal fluorescence detection. A key advantage of LSFM compared to confocal and multiphoton microscopy is that multiple views can be obtained through physical rotation of the specimen that can then be fused together in post processing [3]. The immediate benefits of multi-view imaging and fusion are a more uniform image contrast across the specimen when the sample size is comparable or larger than the scattering length, and a reduction in shadowing artefacts caused by regions within the specimen that absorb excitation light. When combined with deconvolution approaches, multi-view fusion provides a more isotropic and an improved spatial resolution [4–6].

Following the development of conventional two-objective LSFM, the method of oblique plane microscopy (OPM) has been developed to enable LSFM using a single microscope objective to illuminate the specimen and collect the resulting fluorescence [7]. This approach was extended through remote axial scanning of the light-sheet and detection planes, which achieved near-video-rate 3D fluorescence imaging using EMCCD camera technology [8] and 2-colour video-rate 3D fluorescence imaging using sCMOS camera technology [9]. The OPM approach has also been demonstrated for stage-scanned imaging of multi-well plates [10]. Remote lateral scanning of the light-sheet and detection planes was achieved using a rotating polygon mirror [11] and galvo mirrors [12,13]. Folding the remote-refocussing system about a small tilted mirror placed in the focal plane of the second microscope objective [14,15] can be used to increase the numerical aperture of the third microscope objective in the remote-refocussing system [16]. The NA of the third microscope objective can also be increased by using a microscope objective with an NA that approaches unity but with a very small working distance and a front element shaped to allow its close approach to the second microscope objective [17].

While OPM has fewer constraints in terms of sample preparation and the ability to easily image large arrays of specimens in multiwell plates, it does not have the benefits of multi-view LSFM.

In this paper, we present a novel, folded OPM configuration that enables two separate orthogonal views of the specimen to be achieved. Only a single mechanical actuator is required in order to scan the light-sheet and detection plane through the specimen and to switch between the two orthogonal views. This approach enables the benefits of dual-view SPIM to be obtained when performing OPM.

## 2. Methods

### 2.1 Optical setup

The dual-view OPM (dOPM) optical configuration is shown in Fig. 1(a) & (b). For excitation, a multi-wavelength cw laser engine (Omicron LightHUB®) is coupled into a polarisation-maintaining single mode fibre (PMF) and the output is collimated by a 10× air objective (O4, RMS10X, Thorlabs). In order to maximise transmission of linearly polarised excitation light through the folded remote refocussing setup, a visible quarter-wave plate QWP1, (AHWP05M-600, Thorlabs) after O4 is used to generate a circularly polarised beam. The beam is then split into two paths by a visible 50/50 beam-splitter cube (BS, CCM1-BS013/M, Thorlabs). The two beams are then routed by silver mirrors (M1-M5, PFE10-P01, Thorlabs) to generate two light-sheets. In the transmission path of the BS, mirror M1 centres the collimated beam on a cylindrical lens (CY1, 50 mm focal length, LJ1695RM50, Thorlabs). The focal plane of the cylindrical lens is matched to the back focal plane of a 4× air objective lens (O5, RMS4X, Thorlabs) to generate a light-sheet in the front focal plane of O2/3. Mirror M2 then reflects this beam so that it is then at +45° to the optical axis of O2/3, see Fig. 1(b). Similarly, in the reflection path of the BS, steering mirrors M3 & M4, a cylindrical lens (CY2, 50 mm focal length, LJ1695RM50, Thorlabs) and a x4 air objective lens (O6, RMS4X, Thorlabs) generate a light-sheet at −45° to the optical axis of O2/3, see Fig. 1(a).

**Fig. 1.**
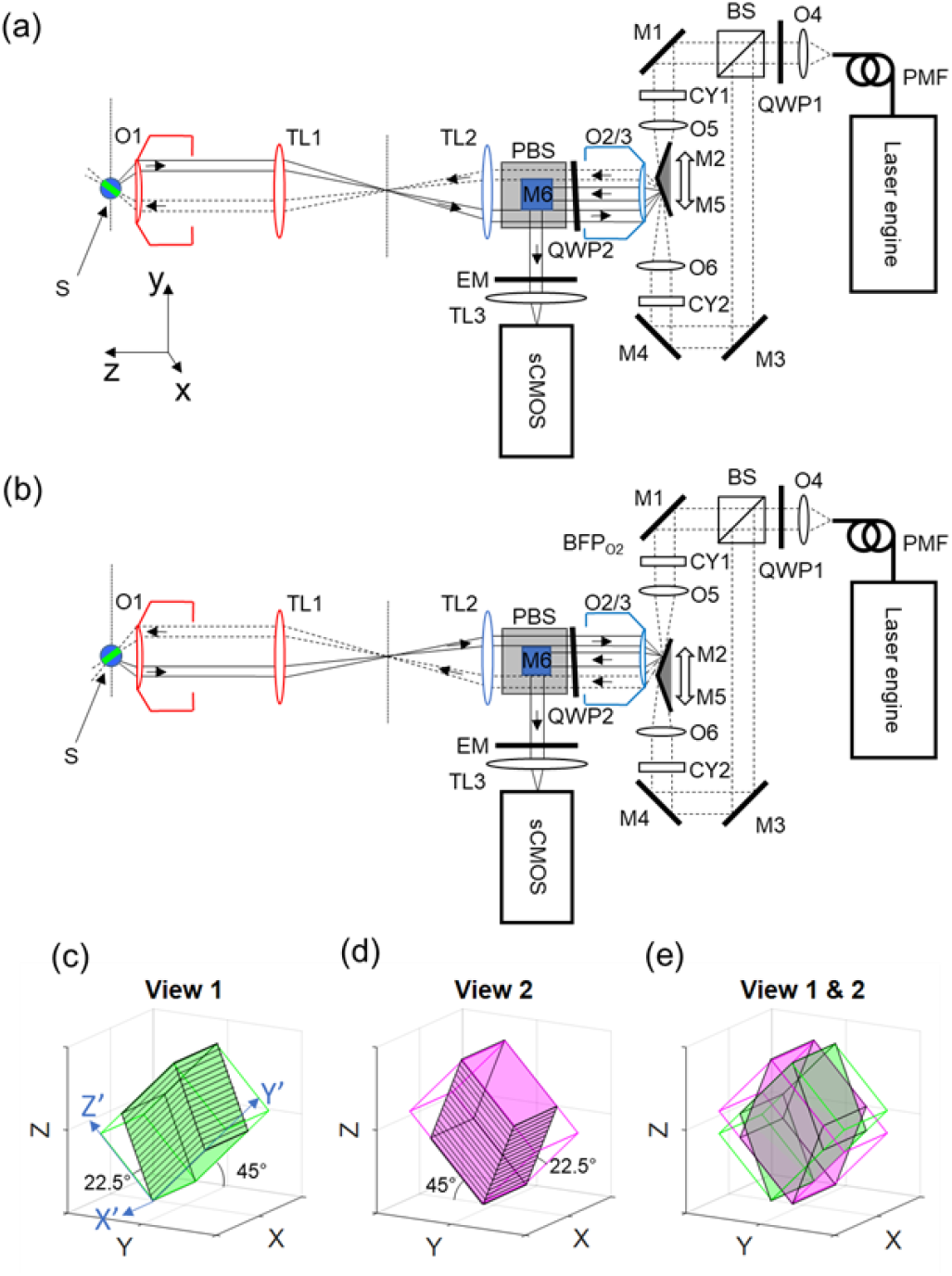
(a) & (b) Dual-view OPM (dOPM) optical configuration, which is based around a remote-refocussing setup folded by mirrors M2 or M5 about the remote microscope objective O2/3. (a) schematic of the optical setup for acquiring View 1. (b) schematic of the system for acquiring View 2. In (a) & (b) O, microscope objective; BFP, back focal plane; TL, tube lens; M, mirror; CY, cylindrical lens; QWP, quarter-wave plate; BS, non-polarising beam splitter; PBS, polarising beam splitter; and EM, emission filter. M2 and M5 are held on a common mount and translated together in the direction shown by the white double-ended arrow. The PBS is oriented so that the reflected fluorescence emission comes vertically up out of the plane of the page and is then reflected into the horizontal plane by M6. The only component that moves between (a) and (b) is the M2 and M5 assembly. The primary microscope is highlighted in red (objective O1 & tube lens TL1) and is a commercially available microscope frame. The secondary microscope is highlighted in blue (objective O2/3 & tube lens TL2). 3D plots (c) and (d) show the outlines of the volumes swept by the two light-sheets. The solid black lines indicate the locations of the acquired image planes as M2 and M5 are scanned and are at ±45° to the optical axis of O1. As M2 and M5 translates, this has the effect of moving the illumination sheet and detection plane at an angle of 22.5° with respect to the illumination/image plane normal. Plot (e) shows the two views superimposed. For plots (c)-(e) the black Cartesian coordinate system corresponds to the primary microscope Cartesian coordinates shown in (a).

As shown by Fig. 1(a) & (b), the light-sheets formed by O5 and O6 are aligned to have their waists coincide with the remote-refocus space of a remote-refocus system formed by the primary (O1 and TL1) and secondary (O2/3 and TL2) microscopes. In this refocus space, a pair of small dielectric coated mirrored prisms held together on a custom mount are used to generate two oblique light-sheets for OPM imaging with two views 90° apart. The mirrored prisms (M2 & M5, MRA03-E02, Thorlabs) are angled with their normal at ±22.5° to the optical axes of O1 and O2/3 to generate light-sheets titled by ±45° relative to the optical axis of the remote-refocusing optics. The folded remote-refocus system is designed for imaging aqueous samples (S in Fig. 1(a)), and the total magnification from S to the remote-refocus space was therefore chosen to be 1.33. The first microscope is based on a Nikon Ti2 Eclipse microscope frame with a 200 mm Nikon tube lens (TL1) and water immersion objective (O1, 40×, Nikon, 1.15 NA, .6 mm WD, MRD77410). The second microscope consists of an air objective (O2/3, 20×, 0.75 NA, Nikon, MRD00205) and a Plössl-style tube lens (TL2, effective focal length 300.8 mm, pair of achromats AC508-1000-A-ML & AC508-400-A-ML, Thorlabs, with achromat spacing optimised in Zemax). A precision #1.5 coverslip (CG15NH1, Thorlabs) is fixed directly to the front surface of O2/3 to account for coverslip correction of O2/3. The mount for mirrors M2 & M5 is fixed on a linear actuator (PIMag®, V 522.1AA & C-413 Motion Controller), with motion direction shown by the white doubled-ended arrow in Fig. 1(a) & (b). This single linear actuator is used to both switch between View 1 (Fig. 1(a)) and View 2 (Fig. 1(b)) and to sweep the light-sheet and plane imaged across the remote refocus image space and therefore across the sample space, S.

The planes illuminated by the two light-sheets are imaged by the folded remote-refocus setup. A second visible quarter wave-plate, QWP2 (AQWP10M-580, Thorlabs) and visible polarising beam splitter (PBS, CCM1 PBS251/M, Thorlabs) allows fluorescence signal from the sample to first transmit through the PBS, be refocused at O2/3 and M2 or M5, reflect vertically from the PBS and be directed horizontally by M6 towards the camera. A pair of emission filters (EM, Semrock, FF03-525/50-25) are used to reject unwanted laser light before fluorescence is imaged by a Plössl-style tube lens, TL3, (effective focal length 325 mm, pair of achromats AC508-750-A-ML & AC508-750-A-ML, Thorlabs) onto a sCMOS camera (ORCA-Fusion, C14440-20UP, Hamamatsu). To achieve the maximum detected fluorescence signal, we orientated the PBS so that it transmits light polarised with its E-field perpendicular to the plane of Fig. 1(a) & (b)). This orientation of the PBS results in the excitation beam being polarised perpendicular to the plane of the figure at the sample. It also results in fluorescence emission that is polarised perpendicular to plane of figure in the sample being transmitted by the PBS on its return from the sample as it propagates towards O2/3. The reflected fluorescence light after its double pass through O2/3 and QWP2 is then polarised in the plane of the figure and so is reflected upwards out of the plane of Fig. 1(a) & (b) and is reflected back to into the plane of the figure by 45° fold mirror M6 placed immediately above the PBS.

In Fig. 1(c) & (d) the 3D plots show the geometry of the volumes scanned in the sample space for the two views. The volume swept in View 1 is shown in green and corresponds to an OPM plane that is rotated anti-clockwise about the x-axis. Similarly, the volume swept in View 2 is shown in magenta and corresponds to an OPM plane that is rotated clockwise about the x axis. It can also be seen that the volumes are parallelepipeds, which is due to the remote refocussing scanning method and is discussed in more detail in section 2.6. Fig. 1(e) shows the two views superimposed for multi-view fusion.

To determine the overall magnification of the OPM microscope, a resolution test target (RES-1, Newport) was imaged. The magnification was determined to be 50.8, corresponding to a pixel size of 0.128 μm in sample space.

### 2.2 Computer hardware

A Tesla-Station Pro-XL workstation (7049GP-TRT, SuperMicro) with Nikon Elements Advance Research software (NIS-Elements) was used to control the microscope acquisition. LabVIEW and DCAM were also used to enable image capture when higher frame rates were required.

### 2.3 Image acquisition of 200 nm fluorescence beads

The NIS-Elements software controlled a DAQ box (USB-6343, NI) to generate TTL signals to synchronise the camera exposure with the laser illumination. The linear actuator controller was configured to respond to an analog voltage from the DAQ box.

The linear actuator controlling the position of M2 and M5 was set to scan ±106 μm along the z axis and about the zero remote refocus positions for each view. When scanning across each view’s volume, the linear actuator was synchronised with the camera in a ‘step and settle’ motion. Following each step, the actuator was configured to settled for 10 ms before each camera image was acquired. The linear actuator position was incremented in steps of 1.22 μm which corresponds to 0.649 μm steps along each views z-axis in sample space (see Section 2.6 for detail on relation between actuator position and sample plane position) resulting in 329 planes per view.

TetraSpeck™ Microspheres, 0.2 μm (T7280, Thermofisher) were imaged with the dOPM system when embedded in agarose gel formed by aqueous agarose (1% agarose) in a glass bottomed dish (35 ml, #1.5 thickness glass bottom dish, MatTek). Compared to the stock solution of beads, the final solution was diluted by a factor of 40. Fluorescence was excited by a 488 nm laser and detected across a 525/50 nm (central wavelength/band pass) band pass emission filter. The average laser excitation power at the sample was 100 μW.

The camera was configured to run in rolling shutter mode with a full frame (2304×2304 pixel, corresponding to a field of view of 295×295 μm^2^ in the sample) and was software triggered. The camera exposure time of 200 ms was synchronised with a flash of the excitation laser, so that the laser was on for the full duration of the camera exposure time. The total acquisition time was approximately 140 seconds. The acquisition speed was limited by software triggering of the camera and the long exposure times compared to the camera readout time of ~10 ms. Higher speeds are possible with hardware triggering and shorter exposure times, see section 2.5.

### 2.4 Image acquisition of 100 nm fluorescence beads

The acquisition was the same as described in detail in following Section (2.5) except lower frame rates of 10 fps and a smaller field of view (1000 × 1000 pixels, corresponding to 128 × 128 μm^2^ in sample space).

TetraSpeck™ Microspheres, 0.1 μm (T7279, Thermofisher) were imaged with the dOPM system when embedded in agarose gel formed by aqueous agarose (1% agarose) in a glass bottomed dish (35 ml, #1.5 thickness glass bottom dish, MatTek). Compared to the stock solution of beads, the final solution was diluted by a factor of 40.

### 2.5 Image acquisition of fixed multi-cellular spheroids

For higher-speed imaging of biological samples, image acquisition was performed during a constant velocity scan of the linear actuator. For this, LabVIEW 2019 was used to control the DAQ box (USB-6343, NI) for hardware-timed control of the laser, linear actuator and camera. The linear actuator controller was configured to respond to an analog voltage input signal from the DAQ box. The linear actuator controller was also configured to output TTL pulses every time the linear actuator moved a predefined distance across two predefined regions of its travel range that corresponded to the scanned volumes of view 1 & 2 (as discussed in Section 2.1). The two TTL output signals associated with each view were combined by a logical OR gate (SN74HC32N, RS-components) and the resulting signal from this OR gate was used to trigger the camera. Specifically, the linear actuator was configured to output TTL pulses for every 1 μm of travel across two 300 μm regions corresponding to views 1 & 2. This corresponded to 300 planes (300 TTL trigger pulses) spaced by 0.532 μm along each view’s z axis direction in sample space (see Section 2.6 for more detail on relation between actuator position and sample plane position). The analog output waveform from from the DAQ box was carefully designed so the linear actuator would linearly ramp across the two 300 μm regions corresponding to view 1 & 2 such that the actuator control box output TTL trigger pulses to produce a frame rate of 90 frames per second (fps). In-between these scan regions, the linear actuator was moved 258 times faster.

A spheroid of 4434 BRAF mutant mouse melanoma cells embedded in Matrigel in one well of a plastic bottomed 96-well plate (PerkinElmer, CellCarrier, #6005550) was imaged with the dOPM system. Actin filaments were fluorescently labelled with Alexa Fluor™ 488 Phalloidin (Thermo Fisher Scientific, Molecular Probes™, #A12379). Fluorescence was excited by a 488 nm laser and detected with a pair of 525/50 nm (centre wavelength/band pass) emission filters.

The camera was controlled by Hamamatsu’s HCImage image acquisition software and was configured to run in rolling shutter mode with a central region of interest of (2000×2000 pixels, corresponding to 256×256 μm^2^ in sample space) and to be externally triggered (ORCA-Fusion, synchronous readout trigger mode) at a frame rate of 90 fps. The total acquisition time for the two volumes totalling 600 planes was 7.06 seconds. The power at the sample plane was 1 mW and this produced an approximately uniform light-sheet across the 550 μm field of view of the objective. The spheroid was approximately 100 μm in diameter, so ~1/5 of the total power was used to illuminate the spheroid.

### 2.6 Image deskewing

To scan the two light-sheets in the microscope sample space, a pair of titled mirrors held on a common mount are positioned in the remote-refocus space to alternately sweep two orthogonal light-sheets. A series of affine transformations describe the corresponding refocussed camera plane position in the sample as it is scanned through one of the two views. Starting in the remote refocus space, the prism mirrors are tilted relative to the x-axis by ±22.5° to rotate the remote refocussed camera plane space by ±45° in the sample. As the linear actuator scans one of the mirrors through the remote-refocus space, the camera plane is refocused along the mirror plane’s normal (±22.5° about the z-axis), see Fig. 1(c) & (d). The refocussing is therefore not normal to the refocussed camera plane. Instead the plane follows a sheared path along the y’-axis according to the amount of z’-axis refocus. This leads to the parallelepiped volume for each view as shown in Fig. 1(c) & (d). For a given translation *d* of the linear actuator in the y direction, the refocus distance in the z direction, *d’* is given by

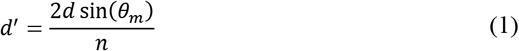

where *θ_m_*=±22.5° is the angle of the mirror normal with respect to the optical axis of the remote refocus system. The amount of refocus normal to the refocused tilted camera plane, i.e. in each views z’ direction is given by

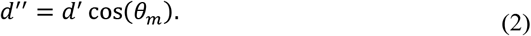

The image plane position depends on the mirror tilt and position in remote refocus space according to translation (T), shear (S) and rotation (R) affine transformations. For View 1 the transformation matrices are,

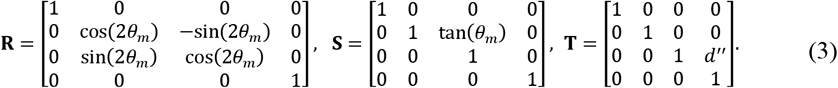

and for View 2 the transformation matrices are,

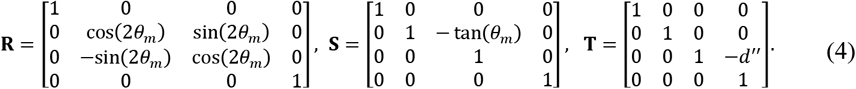

Overall, the coordinates of the imaged OPM plane relative to the camera plane are given by,

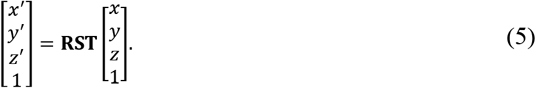

### 2.7 Reconstruction and multi-view fusion using ImageJ

To process the raw data, the Multiview Fusion plugin [18] available in ImageJ was used. The plugin was used to implement the affine transformations described in section 2.6 and an automatic bead-based co-registration procedure for the two views, explained in detail in [18], was used. With the two views co-registered, the same plugin was used to implement interpolation procedures and multi-view fusion and multi-view deconvolution procedures as explained in [5,18]. Deconvolution was performed using the default setting of 10 iterations. For displaying orthoplanes of processed data, the Multiview Fusion plugin was also used to rotate and reslice processed volumes into the microscope Cartesian coordinate frame (as shown in Fig. 1 (a)). Finally, the volumes were exported from the plugin as tiff stacks and the central orthoplanes were extracted.

## 3. Results

### 3.1 Calculation of numerical aperture and fluorescence collection efficiency

By calculating the angular overlap of the angular acceptance cones of O1, O2/3 on the first pass and O2/3 on the second pass [9], the theoretical NA of the dOPM system was determined to be 0.58 in the latitudinal direction and 0.93 in the longitudinal direction, see Fig. 2. This method also provides the fraction of fluorescence emitted isotropically into 2π steradians that would reach the detector in the absence of losses due to the PBS other optical components as 0.16.

**Fig. 2.**
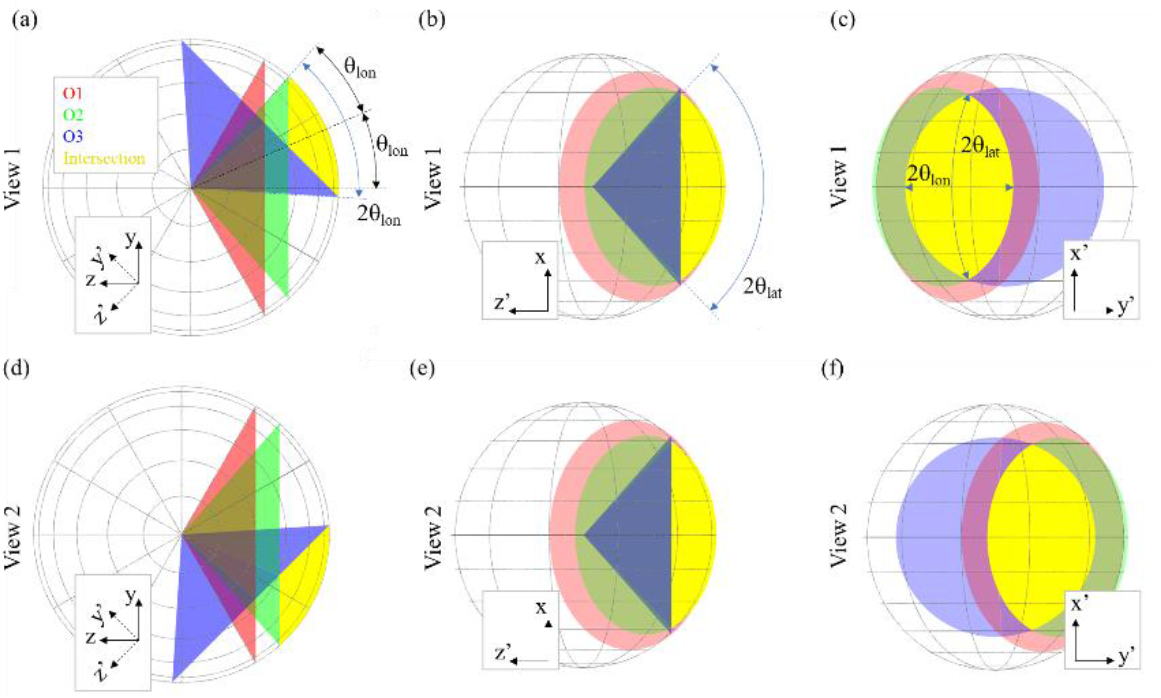
Collection cones of O1, O2, O3 and their intersection for the dOPM system for View 1 shown (a) viewing along the x axis (shown in Fig. 1) and (b) viewing along y axis and (c) viewing along the z axis. The equivalent views are shown for View 2 in (d), (e) and (f). The latitudinal (θ_lat_) and longitudinal (θ_lon_) collection angles of the overall system (intersection) are indicated, together with the tilt (ϕ) of the psf with respect to the optical axis Z in the Z-Y plane.

If the fluorophores in the sample depolarise instantaneously, i.e. have a steady-state fluorescence anisotropy of zero and the fluorescence emission is completely isotropic and unpolarised, then the PBS will transmit 50% of the fluorescence and hence the overall fraction of fluorescence emitted into 2π steradians that reaches the detector is 0.08.

If the fluorophores in the sample have a fixed orientation and do not depolarise, i.e. have a steady state fluorescence anisotropy of 0.4, then the fluorescence emitted is partially polarised [19] and, in the low NA case, then three times the fluorescence signal is polarised perpendicular to the plane of Fig. 1(a) & (b) compared to that which is polarised in the plane of the figure. This assumes that the sample is excited with light polarised perpendicular to plane of figure, as is achieved via the orientation of the PBS shown in Fig. 1(a) & (b). In this scenario, 75% of the fluorescence emitted by the sample reaches the detector in the dOPM setup and hence the overall fraction of fluorescence emitted into 2π steradians that reaches the detector is 0.12. It is important to note that, while the use of the PBS reduces the fluorescence signal reaching the detector, it does not limit the NA of the detection optics.

It can also be seen that the resulting point spread function (PSF) will be tilted in the Z-Y plane by angle ϕ, see Fig. 2(a). The value of ϕ for the configuration used is 18.9°.

### 3.2 Dual-view OPM imaging of beads and characterisation of light-sheet thickness

To quantify the spatial resolution of the dOPM system, a sample of 200 nm fluorescent beads fixed in agarose was imaged. Fig. 3(a) shows a montage of maximum intensity projections for each view and the corresponding 2-colour overlay and fusion when resliced into the Cartesian coordinates of the primary microscope objective (as shown in Fig. 1 (a)) and viewed from the Y-Z perspective. In (a), the green and magenta outlines are used to indicate the outline of the volume scanned by each view. The red line highlights the overlapping region between the two views, which is where information can be combined to improve spatial resolution and contrast. The yellow square highlights the zoomed-in region shown in Fig. 3(b) & (c), which is a maximum intensity projection of the volume.

**Fig. 3.**
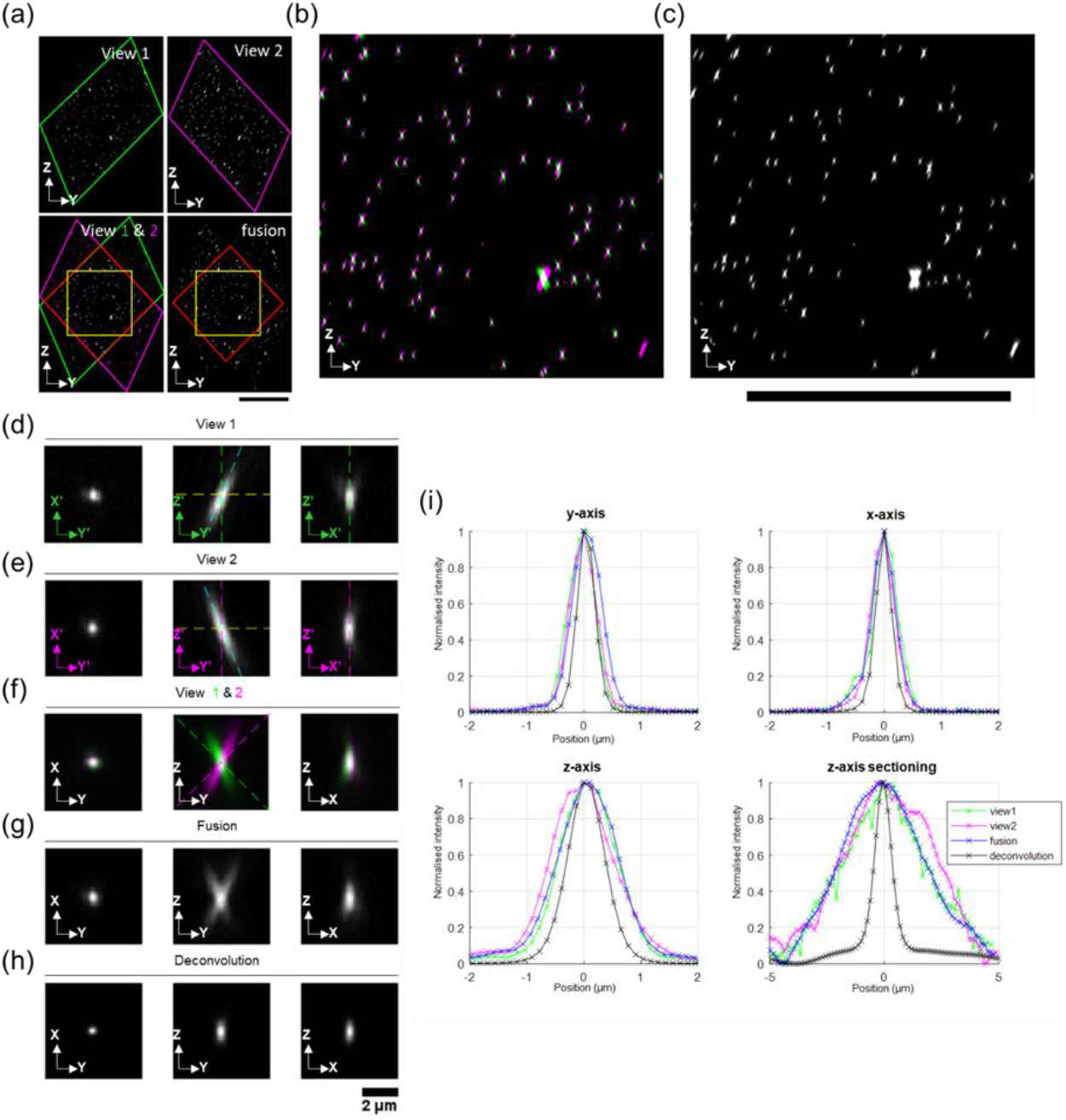
Orthogonal views through dOPM image volumes acquired from 200 nm fluorescent beads embedded in agarose. A 488 nm laser and a 525/50 nm bandpass emission filter were used for fluorescence excitation and detection respectively. In (a), maximum intensity projections for the entire volumes of both views, their 2-colour overlay and their fusion when resliced into the primary microscope Cartesian coordinates (shown in Fig. 1 (a)) are shown viewed from the Y-Z perspective. In (a), the green and magenta outlines show the outline of the volume scanned, and the red line shows the outline of the overlap of the two views. The yellow square shows the zoomed-in region shown in (b) & (c). Figures (d)-(i) show data from a single exemplar bead from the red region highlighted in (a). The montages in (d)-(h) show central orthogonal cuts in the Y-X, Y-Z and X-Z planes for View 1 and View 2, the 2-colour overlays, the two-view fusion and the two-view deconvolution respectively. In (d) & (e), the green and magenta primed coordinates correspond to the Cartesian coordinates for View 1 & View 2 respectively as described in Fig. 1 (c). Also, in (d) & (e), the green and magenta dashed lines show the optical axes (OA) for View 1 & View 2 respectively, which are at ±45° relative to the z-axis of the primary microscope Cartesian coordinates. Also, in (d) & (e), the dashed cyan lines show the tilt ±(45°) of the PSF due to the asymmetric effective detection pupil for each view as detailed in Fig. 2. The montages in (f)-(h) are resliced into the primary microscope Cartesian coordinates. In the 2-colour overlay in (f), the dashed green and magenta lines at ±45° in the Y-Z plane show the optical axes for View 1 & 2 respectively relative to the primary microscope Cartesian coordinates. Plots in (i) show line profiles through the centre of mass of the bead volume fluorescence signal along each axis of the primary microscope Cartesian coordinates. Z-sectioning is reported using the laterally (x,y) integrated signal as a function of depth (along z-axis). The 2 μm scale bar shown below (i) applies to all images across (d)-(h). The scale bar below (a), (b) & (c) is 100 μm.

To show how dOPM combines the information from the two views, Fig. 3(d)-(i) shows data for an exemplar bead taken from the overlapping area of the two volumes shown by the red diamond in Fig. 3(a). Fig. 3(d) shows three central orthogonal cuts taken in the coordinate system of View 1. Fig. 3(e) shows the equivalent images for View 2. Fig. 3(f), (g) & (h) show the 2-colour overlay, the two-view fusion and the two-view deconvolution respectively in the Cartesian coordinate system of the primary objective.

Fig. 3(i) shows central line profiles through the bead’s fluorescence along each axis of the primary microscope Cartesian coordinates. Along each axis, the line profiles are comparable for View 1 & 2 and the fused volumes. For the deconvolved profiles, the x & y profiles are marginally narrower compared to Views 1 & 2 and the fused volume. The line profile in the z direction shows a greater improvement for the deconvolved volume compared to the individual views and fused version. For the z-sectioning (profile obtained by integrating laterally over x and y), there is a pronounced improvement in the deconvolved volume compared to the Views 1 & 2 and the fused volume.

To further quantify and compare the spatial resolution of each view and the fused and deconvolved volumes, Table 1 shows mean full-width half-maximum (FWHM) values for 10 bead image volumes taken from the overlapping area highlighted by the red diamond in Fig. 3(a). The coordinates system used in each row is shown in the left-hand column.

**Table 1.** Mean full-width half-maximum (FWHM) values (and corresponding standard deviation in brackets) for 9 fluorescence bead image volumes from the region within the yellow square shown in Fig. 3. For the z-sectioning, bead signals are laterally integrated (x, y) as a function of depth (z). The Cartesian coordinates correspond to the primary microscope Cartesian coordinates as shown in Fig. 1(a).

| Cartesian coordinate system | Axis | Tilt w.r.t axis | Formula** | Estimated/measured bead FWHM values ($\mu\text{m}$) | | | | |
| --- | --- | --- | --- | --- | --- | --- | --- | --- |
|  |  |  |  | Estimate* | View 1 | View 2 | Fusion | Deconvolution |
| Primary microscope | X | - | $0.51\lambda/\text{NA}_{\text{lon}}$ | 0.37 | 0.46 ($\pm 0.06$) | 0.46 ($\pm 0.02$) | 0.48 ($\pm 0.04$) | 0.35 ($\pm 0.04$) |
| | Y | $18.9^\circ$ | $0.51\lambda/\text{NA}_{\text{lat}}$ | 0.54 | 0.58 ($\pm 0.03$) | 0.59 ($\pm 0.03$) | 0.61 ($\pm 0.04$) | 0.39 ($\pm 0.02$) |
| | Z | $18.9^\circ$ | $1.77\lambda n/\text{NA}_{\text{avg}}^2$ | 2.06 | 1.20 ($\pm 0.07$) | 1.20 ($\pm 0.05$) | 1.39 ($\pm 0.20$) | 0.81 ($\pm 0.07$) |
| | Z sectioning | - | - | - | 5.27 ($\pm 0.66$) | 4.87 ($\pm 0.53$) | 5.17 ($\pm 0.56$) | 1.33 ($\pm 0.45$) |
| View 1 | Z sectioning | - | - | - | 4.99 ($\pm 0.58$) | - | - | - |
| View 2 | Z sectioning | - | - | - | - | 4.89 ($\pm 0.63$) | - | - |

\*Estimated values were calculated using formula shown and include effects of tilt (where applicable), pixel/step size and bead size. \*\* $\lambda$ represents emission wavelength which was 525 nm for the estimate.

As expected, the resolution obtained for each view in the x-direction is better than that achieved in the y-direction due to the higher NA in the latitudinal direction, see Fig. 2. The bead image FWHM measured in the y-direction is broadened by the tilt ϕ=18.9° of the PSF in the coordinate system of the primary objective due to the asymmetric detection pupil shown in Fig. 2.

For both views, the FWHM in the z direction and z-sectioning is worse than the lateral resolution. Compared to each view, the FWHM values for the fused volume are marginally worse. Consistent with Fig. 3(d)-(i), there is a pronounced improvement for the deconvolved beads compared to the individual views and the fused volume.

The expected bead image FWHM were estimated using the scalar formula for the PSF FWHM for a circular pupil, see column 4 of Table 1. The average of the latitudinal and longitudinal NAs was used for the estimate in the axial (z) direction. The tilt of the PSF (if present) was corrected using the appropriate trigonometry, and the pixel/step size and bead size were included by assuming each effect could be modelled as an independent Gaussian distribution. The theoretical FWHM values are reasonably consistent with the measured values and are discussed in more detail in the Discussion section.

The z-sectioning bead image FWHM values given in Table 1 are provided for two different coordinate systems. Those measured in the z-direction of the primary microscope objective enable comparison between the two views, the fused image and the deconvolved image. The z-sectioning values measured in the Cartesian coordinate systems of View 1 and View 2 correspond to the light sheet FWHM in the detection direction of each view and are 4.99±0.58 μm and 4.89±0.63 μm respectively.

To show that dOPM can reveal features that would not be resolvable from a single view alone, the system was applied to image a sample of 100 nm fluorescent beads fixed in agarose that was denser than the sample imaged for Fig. 3. Pairs of beads were identified from the X’-Y’ perspective of each view that were within the measured axial resolution FWHM values for a single view as reported in Table 1. Fig. 4 (a) shows X’-Y’ planes from the perspective of View 1 for each view together with fused and deconvolved versions in the same coordinate system. Fig. 4 (b) shows corresponding line profiles across the pair of beads, which shows that while the two-view fusion marginally improves the ability to resolve the pair of beads by essentially summing the signal from the two views, the two-view deconvolved version makes better use of the lateral resolution information from View 1. A similar trend is shown for a different pair of beads in Fig. 4 (c) & (d) from the View 2 perspective but conversely the better lateral resolution information from View 2 is used to resolve the pair of beads in this X-Y’ plane.

**Fig. 4.**
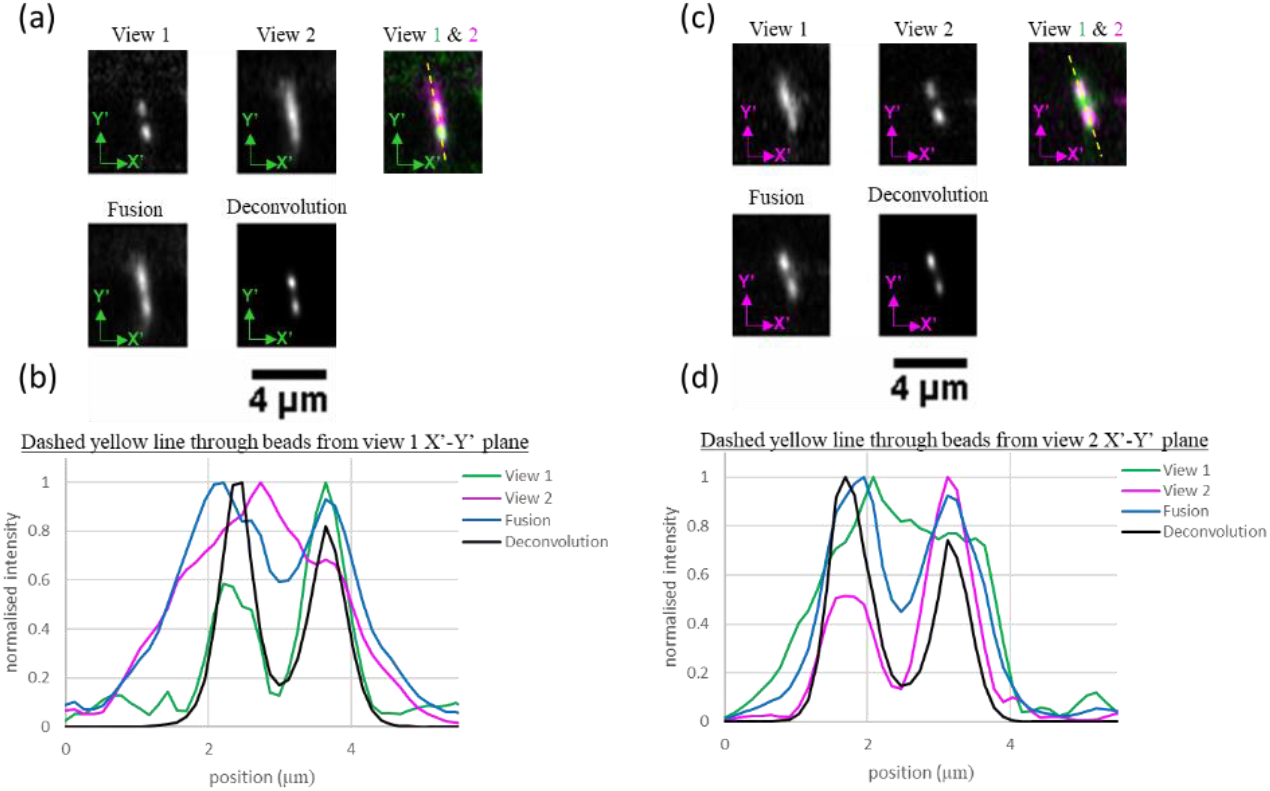
X’-Y’ planes from the perspective of View 1 (a&b) and View 2 (c&d) that intersect pairs of 100 nm fluorescent beads embedded in agarose. A 488 nm laser and a 525/50 nm bandpass emission filter were used for fluorescence excitation and detection respectively. In (a), a montage of common X’-Y’ planes from the View 1 perspective including the two-view fusion and the two-view deconvolution where the green primed coordinates correspond to the Cartesian coordinates for View 1. The 2-colour overlay of data from both views has a dashed line indicating where the line profiles were taken for the plot shown in (b). (c) and (d) show equivalent planes and line profiles using the View 2 perspective for a different pair of beads to those shown in (a) and (b). The magenta primed coordinates correspond to the Cartesian coordinates for View 2. Scale bar in (a) applies to all images as does scale bar in (c).

### 3.3 Dual-view OPM imaging fixed spheroids

The dOPM system was applied to image a fixed multi-cellular spheroid of 4434 BRAF mutant mouse melanoma cells embedded in Matrigel where actin is labelled by Alexa Fluor™ 488 Phalloidin. The spheroid was on the order of 100 μm in diameter. This optically thick, complex biological sample leads to spatial variations in image quality that tend to degrade with optical path length in the sample.

Fig. 5(a)-(f) shows central orthogonal cuts through the acquired image volumes taken with respect to the Cartesian coordinate system of the primary microscope objective (see axes in Fig. 1(a)) for View 1, View 2, the 2 colour overlay (where the images have been registered but not fused), the two-view fused volume and the two-view deconvolved volume. The cartoons in Fig. 5(h)-(j) illustrate the perspectives shown.

**Fig. 5.**
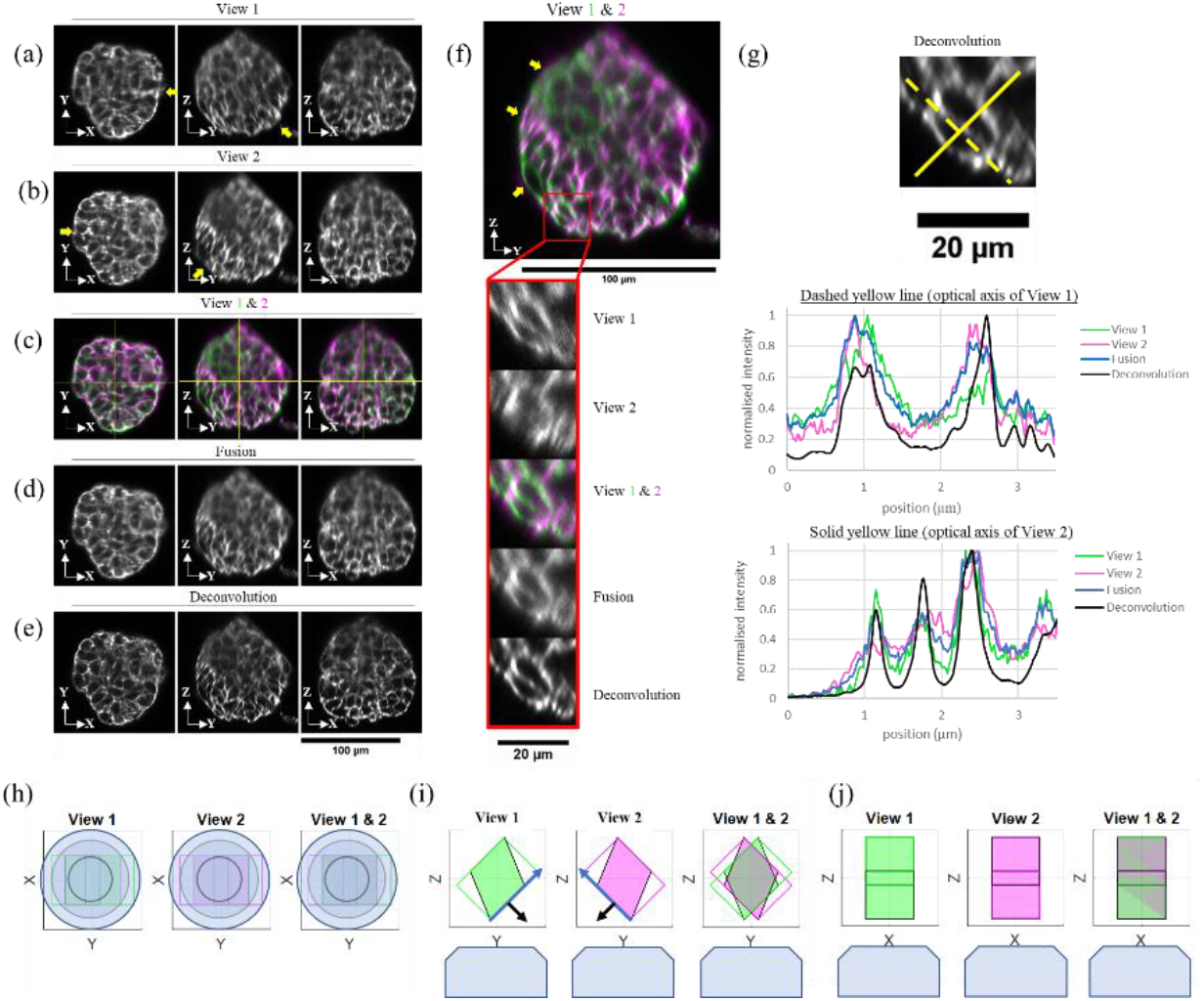
Orthogonal cuts through dOPM image volumes acquired from a fixed spheroid of WMs cells embedded in Matrigel and where Alexa Fluor™ 488 Phalloidin fluorescently labels actin. A 488 nm laser and a 525/50 nm bandpass emission filter were used for fluorescence excitation and detection respectively. In (a), a montage of three images shows central orthogonal cuts in the X-Y, Y-Z and X-Z planes for View 1. (b) shows the equivalent central orthogonal cuts for View 2. In (c), 2-colour overlays are shown for central orthogonal cuts equivalent to (a) and (b) where the View 1 and View 2 volumes have been co-registered but not fused. In (c), yellow lines on the 2-colour overlay images indicate the ortho-plane positions in 3D. In (d), central orthogonal cuts from the two view’s volumes are shown after being co-registered and fused. In (e), central orthogonal cuts from the View 1 and View 2 volumes are shown after being coregistered and deconvolved. In (f), the Y-Z plane shown in (c) is expanded and the region bordered in red shows the location where a subregion has been expanded further. (g) shows line profiles taken from the same sub-region as shown in (f) along the solid and dashed yellow lines indicated at the top of the panel. In (a), (b) & (f), the yellow arrows highlight image features described in the main text. Throughout the figure, the Cartesian coordinate systems corresponds to the Cartesian coordinates of the primary microscope shown in Fig. 1 (a). Scale bar for (a)-(e) is 100 μm. Scale bar for main image in (f) is 100 μm. Scale bar for zoomed region of (f) is 20 μm. In (h)-(j), perspectives plots of the volumes from View 1 and View 2 and their overlay that correspond to the central orthogonal cuts shown in (a)-(f) are shown where View 1 is in green and View 2 is in magenta. In (i), the blue arrow shows the direction of the illumination light-sheet and the black arrow shows the optical axis of the detection optical system for that view.

For View 1, from the central orthogonal X-Y plane shown in Fig. 5(a), the best image contrast is on the right-hand side of the spheroid, whereas for View 2, from the central orthogonal X-Y plane shown in Fig. 5(b), the best image contrast is on the right-hand side of the spheroid (see yellow arrows in Fig. 5(a) & (b)).

For View 1, from the central orthogonal Y-Z plane shown in Fig. 5(a), the best image contrast is on the bottom right of the spheroid, whereas for View 2, from the central orthogonal Z-Y plane shown in Fig. 5(b), the best image contrast is on the bottom left of the spheroid (see yellow arrows in Fig. 5(a) & (b)).

The 2 colour overlay central orthogonal planes shown in Fig. 5(d) and expanded central orthogonal Y-Z plane shown in Fig. 5(f), show that Views 1 and 2 can provide complimentary information, for example, see yellow arrows in Fig. 5(f).

To demonstrate how the two views can be combined to create an improvement spatial contrast across the volume imaged, Fig. 5(d) shows central orthogonal planes from the two-view fused volume that qualitatively are more uniform in image contrast across the central orthogonal planes shown (c.f. Fig. 5(a) & (b)). Similarly, Fig. 5(e) shows central orthogonal planes from the two-view deconvolved volume that, as expected, show qualitatively superior contrast compared to the individual views and the fused view.

To visualise more clearly how the 3D spatial resolution is improved by combining the two views, Fig. 5(f) shows a how a zoomed-in region varies across View 1, View 2, the 2-colour overlay, the fused volume and the deconvolved volume. As shown in the Fig. 5(f), View 1 contributes to the bottom left and top right of the ring-like structures, and View 2 contributes to the top left and bottom right of the ring-like structures.

To illustrate the improvement in information content by combining views, Fig. 5(g) shows line profiles from the same zoomed-in region shown in Fig. 5(f). The dashed yellow line across the zoomed-in region is along the optical axis of View 1 and shows two peaks and a trough in intensity that are contrasted better by View 2 as expected. Conversely, the solid yellow line across the zoomed-in region is along the optical axis of View 2 and shows three peaks and two troughs in intensity that are contrasted better by View 1 as expected.

## 4. Discussion

In section 3.2 we measured the size of images of 200 nm fluorescent beads and compared the results to the values expected from the scalar theory accounting for bead size, camera pixel size and tilt of the PSF where relevant. In the y-direction of the primary microscope objective – which corresponds to the longitudinal NA of the system – the experimental values obtained for View 1 and View 2 match the estimated values to within 9%. In the x-direction of the primary microscope objective – corresponding to the latitudinal NA of the system – the corresponding values agree to within 24%. This poorer agreement is attributed to the higher NA of the system in this direction, which increases the importance of fluorescence anisotropy effects and reduces the validity of the scalar PSF estimate. A more exact theoretical calculation will be carried out in the future. We note that this has been calculated previously for a folded-OPM system [16], but that this analysis did not fully include the effect of the tilted fold mirror.

The NA and fluorescence collection efficiency of the dOPM implementation reported here are limited by the NA of microscope objective O2. In the future, these can be increased without compromising the field of view by instead using e.g. an Olympus 20x/0.8 lens. Increasing the NA of O1, e.g. to a 60x/1.2 water immersion lens or a 60x/1.27 water immersion lens, coupled with the use of a 50x/0.95 lens for O2 would further increase the collection efficiency and spatial resolution achieved at the expense of decreased field of view and working distance of O1. The calculated NAs and collection efficiencies for some different potential dOPM microscope objective combinations are summarised in Table 2. We note that the collection efficiency for the 60x/1.2W configuration shown in Table 2 has a collection efficiency of 0.22 in the case of a steady-state anisotropy of zero, which is higher than the collection efficiency for the equivalent previously published OPM configuration employing a 40x/0.6 lens for the third microscope objective [9] despite the fact that half of the signal is lost at the PBS in dOPM.

**Table 2.** Summary of NA and collection efficiencies of fluorescence emitted into 2π steradians (CE) for different choices of dOPM microscope objectives O1 and O2.

|  |  |  |  |  |
| --- | --- | --- | --- | --- |
| O1 | 40x/1.15W | 40x/1.15W | 60x/1.2W | 60x/1.27W |
| O2 | 20x/0.75 | 20/0.8 | 50x/0.95 | 50x/0.95 |
| $\theta_{\text{OPM}}$ | 45° | 45° | 35° | 25° |
| $\text{NA}_{\text{lat}}$ | 0.93 | 1.01 | 1.03 | 1.14 |
| $\text{NA}_{\text{lon}}$ | 0.58 | 0.68 | 1.20 | 1.26 |
| CE for O1 | 0.50 | 0.50 | 0.57 | 0.70 |
| dOPM CE* for steady-state anisotropy = 0 | 0.08 | 0.10 | 0.22 | 0.28 |
| dOPM CE* for steady-state anisotropy = 0.4 | 0.12 | 0.15 | 0.33 | 0.42 |

\*CE including loss at PBS.

dOPM has a number of advantages. First, it allows two views of the sample to be obtained whilst requiring only two microscope objectives in the remote-refocussing setup, which reduces cost compared to OPM. Similarly, only one computer-controlled actuator is required to achieve both switching between views and for scanning during acquisition of each view, further reducing cost. The actuator used in this demonstration can perform saw-tooth operation at 25 Hz, and so the dOPM configuration also has the potential to achieve video-rate volumetric imaging. Furthermore, by scanning the illumination and detection plane by scanning the M2 & M5 assembly means that the mass scanned is low compared to that of a microscope objective in the case of OPM [8] and doesn’t require an additional 4-f system to allow a galvo mirror to be placed conjugate to the pupil planes of O1 and O2 as required for SCAPE [12,13] and SOPi [12]. Finally, the folded remote-refocussing geometry allows the numerical aperture of the 3rd microscope objective in the remote-refocussing optics to have the same numerical aperture of the 2nd microscope objective.

## 5. Conclusion

We have demonstrated a new OPM geometry capable of acquiring two orthogonal views of the sample that can then be fused in post-processing to reduce sample-induced image artefacts. Furthermore, two-view deconvolution can be implemented to obtain a more isotropic PSF and to spatially resolve features not possible with a single view alone. Using a water immersion 40× 1.15 NA primary objective and two views at ±45°, we measured the FWHM of deconvolved image volumes of 200 nm fluorescent beads to be 0.35±0.04 μm, 0.39±0.02 μm and 0.81±0.07 μm in the x, y and z -directions respectively. The laterally integrated z-sectioning value was 1.33±0.45 μm. This was achieved with light-sheet FWHM in the frames of the two views of 4.99±0.58 μm and 4.89±0.63 μm. We also demonstrated the performance of the system for imaging a ~100 μm diameter spheroid. The fusion and deconvolution of the two views reduced inhomogeneities due to shadow artefacts and provided a more uniform resolution around a ring-shaped feature. A single computer-controlled actuator is employed to both switch between the two views and to scan the light-sheet and detection planes through the sample when acquiring each view. The use of a folded remote-refocussing setup is compact and only uses two microscope objectives but causes some light loss due to the need for the PBS. However, the folded OPM configuration does have the advantage that the NA of the third microscope objective is the same as that of the second objective, as the small tilted fold mirror allows for a greater tilt angle than could be achieved when using identical lenses for the second and third objectives in a non-folded configuration.

## Acknowledgements

The authors would like to acknowledge the expert help from Martin Kehoe, Simon Johnson and John Murphy in the Optics Workshop of the Photonics Group of Imperial College London who helped design and fabricate the components for the light-sheet microscope system.

